# Ovarian cancer G protein-coupled receptor 1 inhibits A549 cells Migration through Casein kinase 2α intronless gene

**DOI:** 10.1101/556720

**Authors:** Adhikarimayum Lakhikumar Sharma, Puyam Milan Meitei, Naorem Tarundas Singh, Thiyam Ramsing Singh, Lisam Shanjukumar Singh

## Abstract

We have previously reported that ovarian cancer G protein-coupled receptor 1 (OGR1) is a new metastasis suppressor gene. We have also reported for the first time that a new intronless gene for casein kinase 2α (*CSNK2A3*) is expressed in human cells. The promoter of the well-known casein kinase 2α (*CSNK2A1*) displays characteristics of housekeeping gene whereas *CSNK2A3* has a characteristic of a regulated promoter with two TATA boxes and a CAAT box. In this study, we found that OGR1 up-regulates expression of *CSNK2A3* by about 3 folds in A549 cells but not *CSNK2A1*. OGR1 also up-regulates expression of neutral endopeptidase (NEP). The OGR1 induced inhibition of A549 cell migration is completely abrogated by inhibition of casein kinase 2α activity, whereas partial abrogation (~ 30%) was observed in the presence of NEP inhibition. The results also revealed that OGR1 regulates *CSNK2A3* via activation of Rac1/cdc42 and MAPKs pathways. CK2 is ubiquitously expressed and in contrast it is believed to be a constitutively active enzyme and its regulation appears to be independent of known second messengers. There is no previous report on how expression of CK2α in cancer cells is regulated although many studies have report of aberrant expression of the kinase in cancer. In the current study, we are reporting for the first time the regulation of intronless casein kinase 2α gene, *CSNK2A3* in cancer cells. Our findings suggest that the aberrantly casein kinase 2α expression found in various cancer cells may the due to *CSNK2A3* expression which is potentially regulated by several master regulators of the developmental pathways rather than well-known casein kinase 2α gene, *CSNK2A1*.

## Introduction

The ovarian cancer G protein-coupled receptor 1 (OGR1) and related subfamily members mediate the functions of several lysophospholipids which include endothelial barrier function, endothelial cell proliferation, migration, and tube formation, T cell migration, glucocorticoid-induced thymocyte apoptosis, and globoid cell formation [1–5]. OGR1 gene has been shown to be expressed at lower levels in metastatic compared with primary prostate cancer tissues [6]. Recently, we have shown using an orthotopic mouse metastasis model that OGR1 is a novel metastasis suppressor gene for prostate cancer when it is re-expressed in cancer cells [5]. Further, other researchers have also reported that OGR1 inhibits breast and ovarian cancer cells *in vitro* when it is re-expressed in cancer cells [7, 8]. However, our previous study revealed that OGR1 has inhibitory effect on prostate cancer tumorigenesis when expressed in host/stromal cells using OGR1 knockout TRAMP mice prostate cancer model [9]. Further, we have also demonstrated that OGR1 significantly inhibited melanoma tumorigenesis in OGR1 knockout mice [10]. Another study of Horman SR, *et al*. has reported that murine colon tumor implants in OGR1 knockout mice displayed delayed tumor growth [11]. Therefore, these previous reports clearly indicate that in contrast to OGR1’s tumor-suppressing role in tumor cells, host cell OGR1 may be involved in and/or required for tumor growth. OGR1 and its subfamily G protein-coupled receptors (GPCRs), GPR4, G2A and T-cell death-associated gene 8 (TDAG8), have been shown to have proton-sensing ability [12, 13]. Many of these proton sensing activities have been identified in cells over-expressing one or more of these GPCRs. More recently, proton sensing activities have been detected in cells from GPR4- and TDAG8-, but not G2A-deficient mice [3, 14, 15]. Previous studies have reported that OGR1 activity on tumorigenesis is independent of its pH-sensing activities. These G-protein coupled receptors also appear to have ligand-independent constitutive activity [5, 7, 8, 10]. However, the signalling pathway of OGR1 in inhibition of metastasis is not clearly understood. One previous report has revealed that OGR1 induced activation of Rho but down-regulation of Rac1 in breast cancer cell line [8]. Therefore, it is important to investigate the down-stream cellular proteins of OGR1 in its function as metastasis suppressor gene.

Casein kinase 2α (CK2α) acts as a regulator of several hallmarks of cancer cell behaviour [16–18]. CK2α gene may potentially be induced or repressed by several master regulators of developmental pathways [19–21]. We have for the first time reported that an intronless gene of CK2α (*CSNK2A3*) is expressed in human megakaryocytic cells [21]. Recently other studies have revealed that *CSNK2A3* is expressed in 293T, A549 and NIH-3T3 cells and further polymorphism of *CSNK2A3* plays oncogenic roles in lung cancer [22]. Importantly, although the activity of the CSNK2A1 gene has been shown to be elevated in human cancers, no solid genetic or epigenetic evidence is available regarding the cause of the high-activity of CK2α in cancer cells. CK2 proteins are upregulated in the human tumors tested so far, suggesting its important role in cancer progression [23, 24]. In cancer, CK2 is proposed to regulate essential cellular processes such as cell growth [25], cell proliferation [26], cell survival [27], cell morphology [28], cell transformation [29, 30] and angiogenesis [31]. Although, it is observed that there are multiple layers of regulation of CK2α expression [24, 32], reports on regulation of CK2α expression is very limited. The original view in the literature is that CK2 is predominantly regulated post-transcriptionally; however, recent studies strongly suggest that regulation at the transcriptional level is also important in some cancers [33]. Recently, Das, N et al have reported that ERα transcriptionally activates CKα [34]. Unpredictably, some cancers show under-expression of CK2 transcripts in breast, ovarian, and pancreatic cancer [33]. Furthermore, CK2 transcript levels could have a prognostic value in cancers (e.g. CK2α in squamous cell carcinoma of the lung). For the most part, high levels of CK2 transcript correlate with lower overall survival (e.g. breast and ovarian cancer, glioblastoma, kidney and liver cancer) [33, 35–37]. However, in lung adenocarcinoma, high levels of *CSNK2A2* (CK2α’) and *CSNK2A3* correlate with higher survival rates [24, 33]. Similar to lung adenocarcinoma, overexpression of *CSNK2A3* in renal clear cell carcinoma led to increased survival [24]. From the above data, it is not clear whether CK2 is anti-cancer or pro-cancer molecule.

Neutral endopeptidase 24.11 (NEP, neprilysin, enkephalinase, CD 10) is a widely distributed membrane enzyme, occurring on a variety of cells [38]. The biological and regulatory effects of NEP are presumed only to result from its enzymatic function [39, 40]. However, recent data suggest that NEP protein expression in of itself can effect signal transduction pathways that regulate cell growth [41, 42] and apoptosis [43]. Tokuhara et al reported tumors with high NEP and low CD13 were associated with better prognoses [44]. Gurel et al reported that in lung squamous cell carcinomas both tumoral and stromal NEP expression were unfavorable prognostic factors, while in non-squamous cell carcinomas tumoral NEP expression was a favorable prognostic factor [45]. In a study conducted by Kristiansen et al, tumoral NEP expression was not associated with prognosis in NSCLCs [46]. Ono et al reported that neither tumoral nor stromal NEP expression correlated with prognosis in stage I lung squamous cell carcinomas [47]. Recently, Leithner et al reported that high NEP expression, based on gene expression analysis, was associated with unfavorable prognoses in lung adenocarcinomas but not in non-adenocarcinomas [48]. However, it is unclear whether this result was based on tumoral or stromal NEP expression because localization of NEP was not confirmed via immunohistochemistry. Therefore, the prognostic value of tumoral NEP for early-stage lung adenocarcinoma remains unknown. NEP expression was lower in certain carcinomas of the lung than in adjacent normal tissue [38]. In the current study we aim to identify the key cellular protein(s) involved in the inhibition cell migratory induced by a metastasis suppressor gene, OGR1.

## Results

### OGR1 regulates expression of CSNK2A3 and NEP but not CSNK2A1

To investigate the downstream molecules involved in the OGR1 induced inhibition of cancer cells migration, OGR1 was over-expressed transiently in A549 cells for 48 hrs and transcript expression of *CSNK2A1*, *CSNK2A3* and *NEP* were analysed using semi-qPCR. Initially, OGR1 over-expression in A549 was confirmed by semi-qPCR. The results showed that OGR1 strongly up-regulates transcript expression of *CSNK2A3* (CK2α intronless gene, CK2αP) and NEP. However, OGR1 does not affect significantly the expression of *CSNK2A1* in A549 cells (**Figure 1 A, B**). Further, protein expression of CK2α and NEP were analysed by immunoblotting upon OGR1 over-expression in A549 cells using respective specific antibodies (**Figure 1C, D**). The over-expression of OGR1 protein was confirmed using specific antibody against OGR1. The specific antibody for CK2α recognises proteins of both *CSNK2A1* and *CSNK2A3* since there are only four amino acids different in the sequence of the two genes [21]. Taking together the results of transcript expression of casein kinase 2α genes (CSNK2A1 and *CSNK2A3*) and protein expression of CK2α, it is clearly indicated that the increased in protein of CK2α in immunoblotting may be the product of *CSNK2A3* (*CK2αP*). The immunoblotting against NEP confirmed that OGR1 over-expression cells increases NEP expression in A549 cells (**Figure 1C**). The overall results indicate that OGR1 up-regulates expression of *CSNK2A3* and *NEP*, but not *CSNK2A1* in lung cancer cells. Our finding is the first report to demonstrate that the expression of *CSNK2A3* is regulated. There is very limited report of how CK2α gene expression is regulated inspite of many reports on aberrantly expression in cancer.

**Figure 1:**
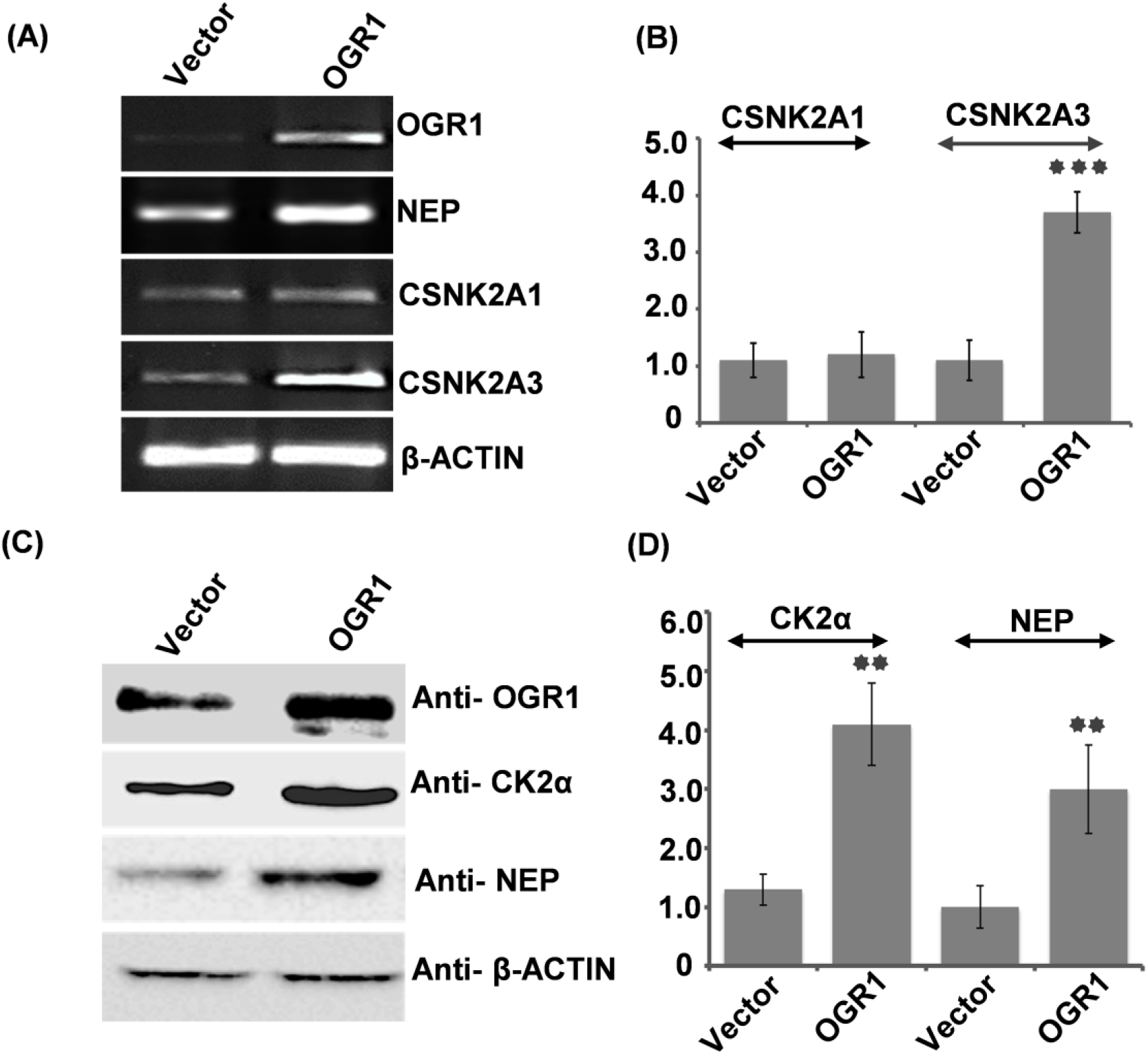
OGR1 up-regulates CSNK2A3 (CK2α intronless gene) and NEP but not CSNK2A1 in A549. (**A**) A549 cells were transfected with OGR1 (OGR1) or pcDNA3.1 (Vector), transcript expressions of *CSNK2A1, CSNK2A3* and *NEP* were analysed in the by semi-quantitative RT-PCR. β-actin was used as control for equal loading. (**B**) RNA bands intensities for *CSNK2A1* and *CSNK2A3* were analysed by using Image Acquisition and Analysis software (BioRad, USA) and presented in graph. (**C)** Protein expressions of NEP and CK2α were assayed by Western blotting with specific antibodies and β-actin was used as control for equal landing, (**D**) protein band intensities were analysed by using ‒a Image Acquisition and Analysis software (BioRad, USA) and presented as graph. Experiments were repeated thrice. Statistical significance between control (Vector) and OGR1 was calculated using student’s “t” test. Bars indicates SD, ** indicates p-value < 0.01 and *** indicates p-value <0.001.

### CK2αP is up stream of NEP in the OGR1 signalling pathway

To investigate whether CK2αP is upstream of NEP or vice-versa in the OGR1 signalling pathway, we assessed expression of both the proteins in the presence of either specific chemical inhibitor of CK2, CX-4945 and immunoblotting against NEP or specific chemical inhibitor of NEP, thiophan and immunoblotting against CK2α after A549 cells were transiently transfected with OGR1 or empty vector (control). Interestingly, the results showed that in the presence of CX-4945, increased expression of NEP induced by OGR1 is abrogated to a similar level of control cells (**Figure 2A, B**). However, thiorphan did not affect expression of CK2α protein in A549 cells (**Figure 2C, D**). These results indicated that CK2αP is upstream of NEP in the signalling pathway of OGR1. It is previously reported that CK2α regulates NEP activity by phosphorylating at cytoplasmic tail [49].

**Figure 2:**
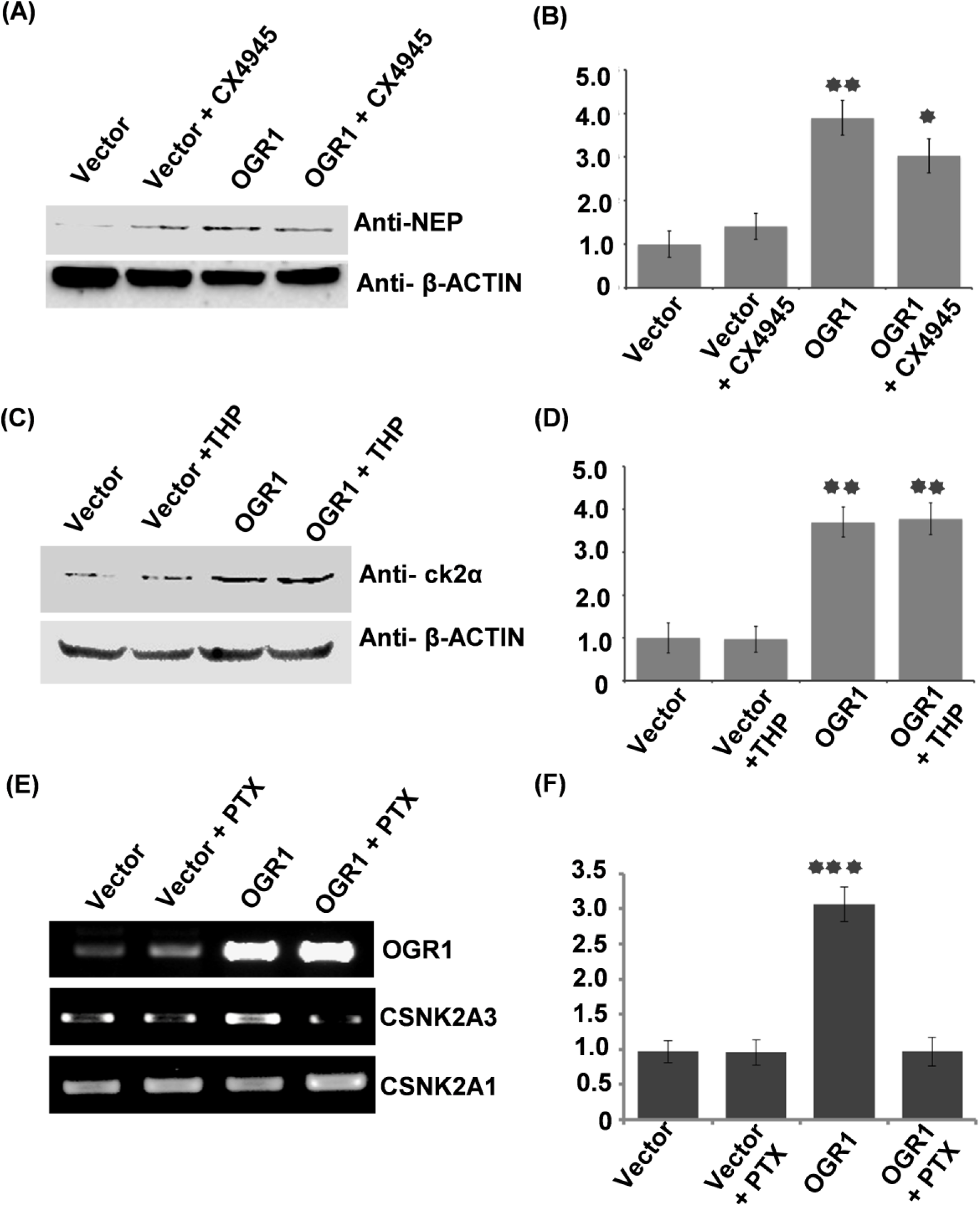
CK2 is upstream of NEP in the OGR1 signalling pathway. Western blot assay of NEP and CK2α expression in presence or absence of specific inhibitors. (**A**) A549 cells were transfected with pDNA3.1-OGR1 (OGR1) or pcDNA3.1 (Vector) and NEP expression was analysed with anti NEP in presence or absence of CX-4945, CK2 inhibitor (CX4945) and **(B)** protein band intensities were analysed by using Image Acquisition and Analysis software (BioRad, USA) and presented as graph, (**C**) expression of CK2α was analysed with anti CK2 in presence or absence of NEP inhibitor, thiophan (THP) and (**D**) protein band intensities were analysed by using Image Acquisition and Analysis software (BioRad, USA) and presented as graph, (**E**) Semi Quantitaive PCR showing the expression of NEP and CK2 in presence or absence of Gi inhibitor PTX and (**F**) intensities of protein bands were analysed by using Image Acquisition and Analysis software (BioRad, USA) and presented in graph. Stastatical significance between treated and untreated was calculated using student’s “t” test. Bars indicate SD, * indicates p-value < 0.04, ** indicates p-value < 0.01 and *** indicates p-value <0.001.

Further, we have reported earlier that OGR1 inhibits PC3 cell migration via Gαi activation [5]. Therefore, to investigate whether OGR1 induced up-regulation of *CSNK2A3* in A549 is dependent of activation of Gαi, we analysed the transcript expression of *CSNK2A3* in the presence or absence of pertussis toxin (PTX) upon OGR1 over-expression or empty vector. We used transcript expression of CSNK2A1 as control. The result revealed that the presence of PTX abrogated the OGR1 induced up-regulation of expression of *CSNK2A3* (**Figure 2E, F**).

### CK2αP functionally involve in OGR1 induced inhibition of lung cancer migration

We have previously demonstrated that OGR1 is a metastatic suppressor gene *in vitro* as well as *in vivo* using an orthotopic mouse metastasis model [5]. In the current study, we found that OGR1 induces increased expression of CK2αP and NEP in transcript and protein levels. Therefore, in the current study, we investigate functional roles of both CK2α and NEP in the OGR1 induced inhibition of A549 cell migration *in vitro* using wound healing model in the presence or absence of inhibitor of CK2α and NEP. The results showed that OGR1 inhibits migration of A549 cells. In the presence of CK2 specific inhibitor, CX-4945, the OGR1 induced inhibition of A549 cells migration was completely abrogated (**Figure 3A, B**). However, thiophan decreases the OGR1 induced inhibition of A549 cell migration only by ~30%, indicating there may be other molecule(s) other than NEP through which CK2αP induces inhibition of cancer cell migration following the action of OGR1.

**Figure 3:**
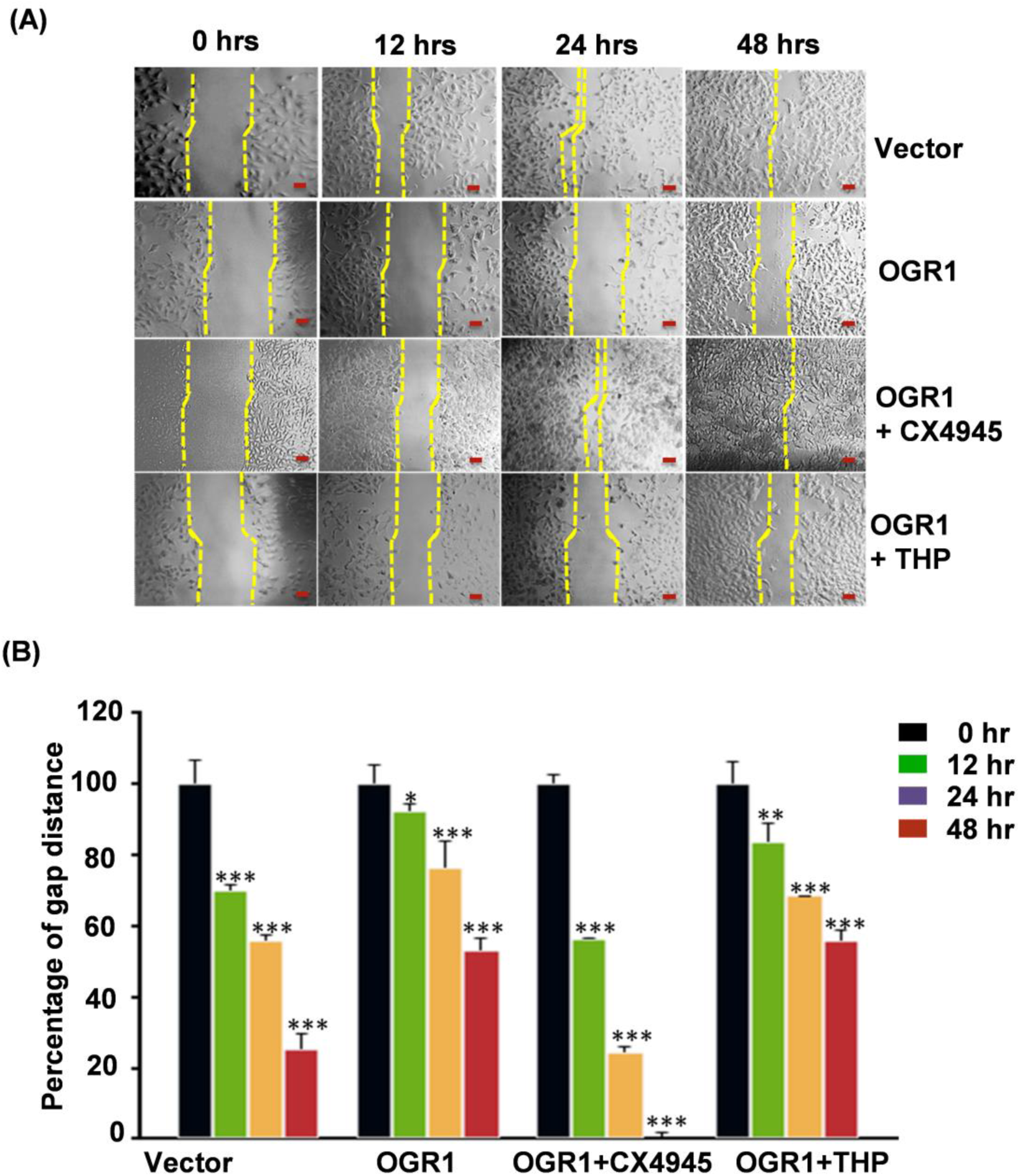
OGR1 inhibits A549 cell migration via CK2. (**A**) OGR1 inhibits A549 cell migration but presence of CK2 inhibitor, CX4945 abrogated the effect of OGR1 but not the presence of NEP inhibitor, thiophan (THP). A549 cell were transfected with pDNA3.1-OGR1 or pcDNA3.1 (Vector) in presence of CK2 and NEP inhibitors. Cell migration was analysed and image was taken hourly. (**B**) To measure the area of wound, the length of scratches were measured on the light microscope and plotted on graph. Wound area of control 0 hr was taken as 100% and means & SD. P-values were calculated comparing with the control values of the same hrs of treatment by students’ “t” test. The bar (Red) indicates the scale of 100µm. Bars indicate SD, * indicates p-value < 0.04, ** indicates p-value < 0.01 and *** indicates p-value <0.001.

### Involvement of Rac1 and cdc42

The known role of the small GTPase; Rac1 and cdc42 as major drivers of cell motility prompted us to assess whether Rac1 and cdc42 are involved in the OGR1 induced up-regulation of CK2αP expression which leads to inhibition of A549 cells migration. A549 cells were transiently transfected with OGR1 alone or co-transfected with cdcT17N (dominant negative of cdc42) or RacT17N (dominant negative of Rac1). Similarly, control A549 cells were also transiently transfected with empty vector alone and co-transfected with cdcT17N or RacT17N. Initially, over-expression of OGR1 in the OGR1 transfected cells was confirmed by semi-qPCR. The results showed that in the presence of cdcT17N and RacT17N, the OGR1 induced up-regulation of CK2αP in A549 cells was significantly abrogated, suggesting that both small G proteins cdc42 and Rac1 are involved in the OGR1 induced up-regulation of CK2αP in A549 cells (**Figure 4A**).

**Figure 4:**
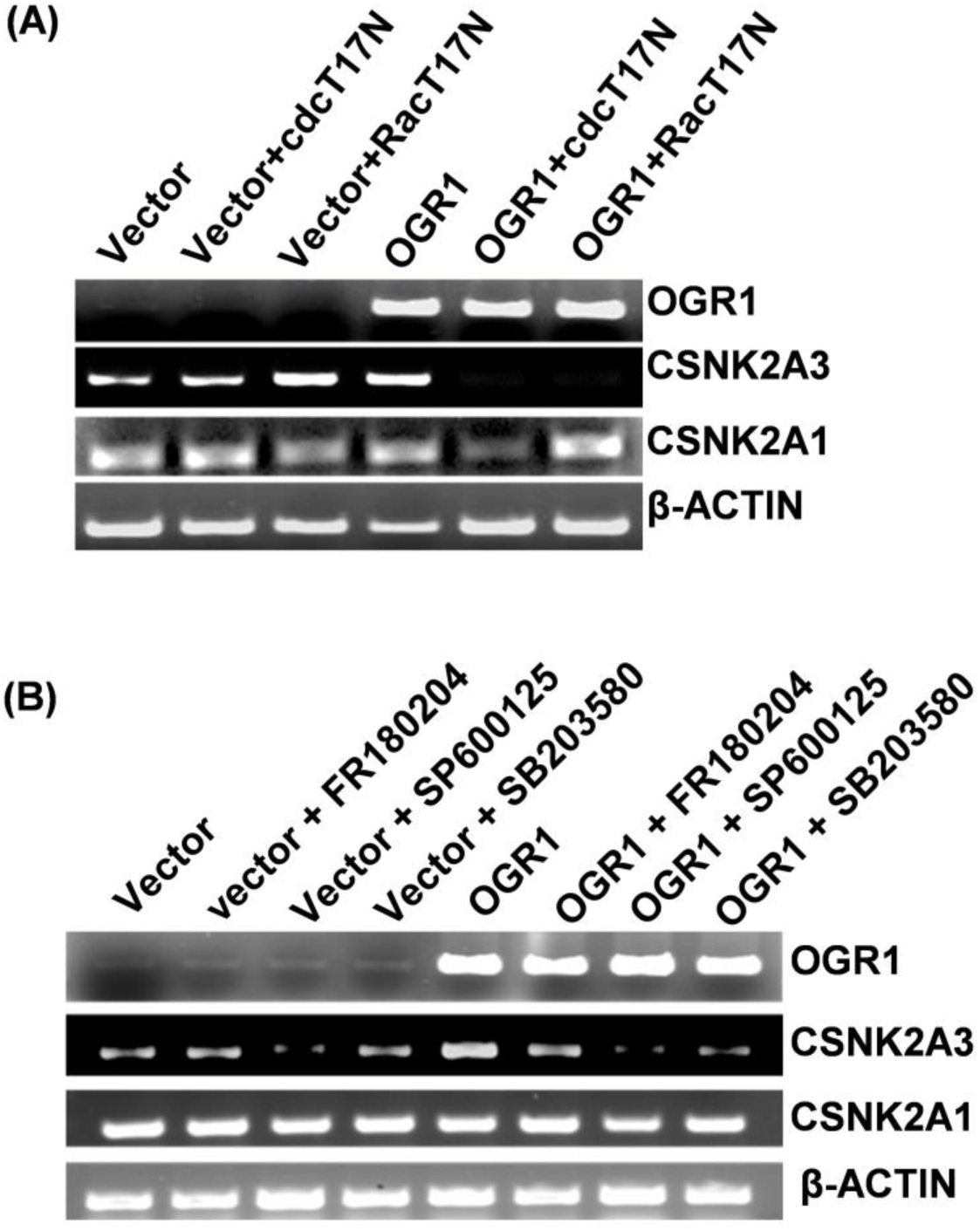
Role of Rac/CDC42 and MAPK pathways in regulation of CK2α genes induced by OGR1. **(A)** A549 cells were co-transfected with OGR1 or pcDNA3.1 (Vector) and negative dominant mutants of Rac (pcDNA3.1-RacT17N) or negative dominant mutants of CDC42 (pcDNA3.1-cdcT17N). Semiquantitative RT-PCR of was performed to analyse *CSNK2A1* and *CSNK2A3* transcript expression. β-actin was used as control for equal loading; Inhibition of Rac activity abrogated up-regulation of *CSNK2A3* induced by OGR1 **(B)** A549 cells were also transfected with pDNA3.1-OGR1 or pcDNA3.1 (Vector) in presence of MAPK kinase inhibitors. Semi-quantitative RT-PCR of was performed to analyse *CSNK2A1* and *CSNK2A3* transcript expression. β-actin was used as control for equal loading; inhibition of JNK and p38 using specific inhibitors, SP600125 and SB2203580 respectively, abrogated the up-regulate of *CSNK2A3* induced by OGR1 but not by inhibition of ERK (FR180204) whereas inhibition of ERK, JNK and p38 does not significantly affect the expression of *CSNK2A1* in A549.

**Figure 5.**
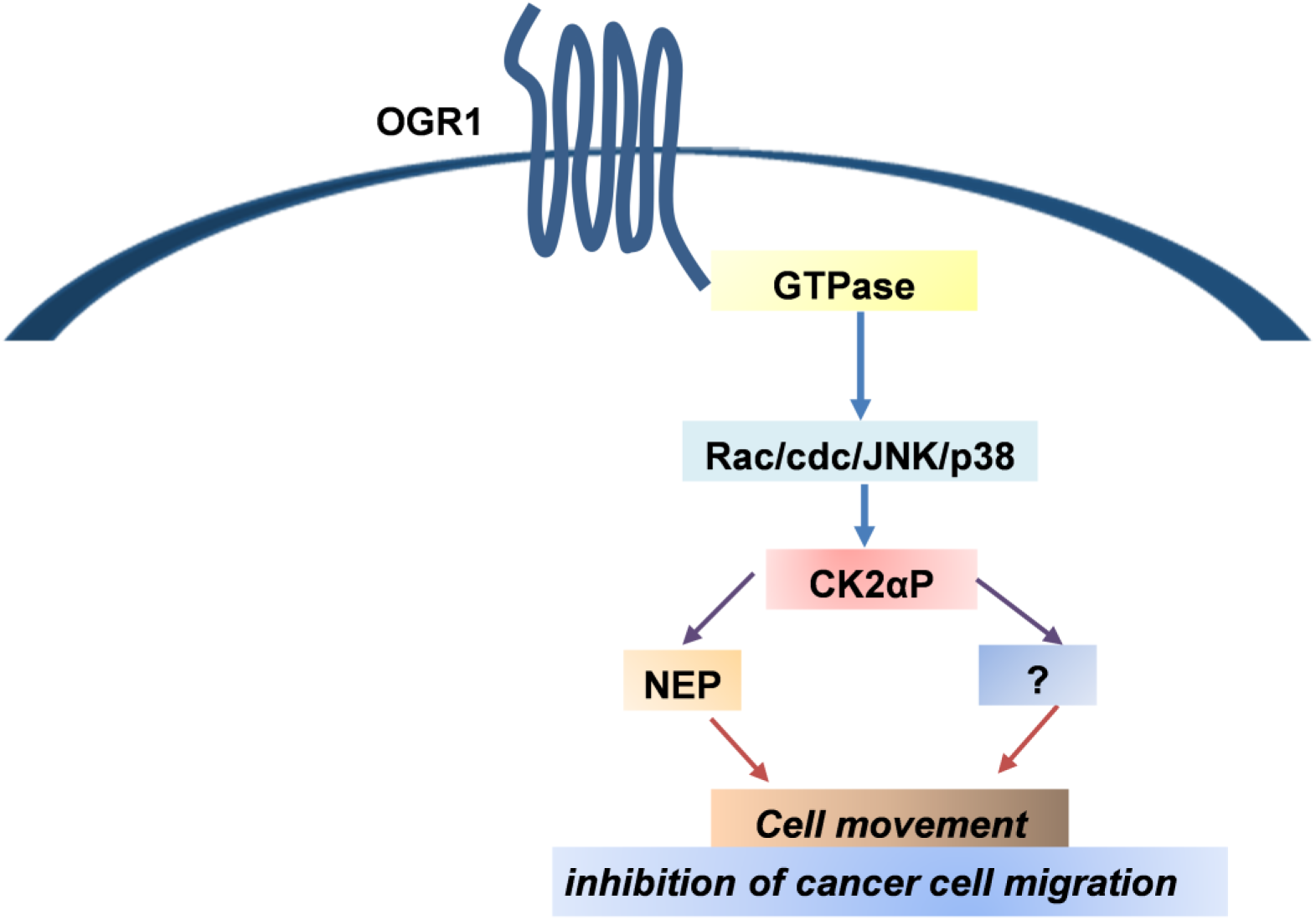
Speculative signaling pathways of OGR1 for inhibition of A549 cell migration is illustrated. OGR1 activates protein Gαi which leads to further activation of small G proteins Rac1 and cdc42. Activations of Rac1 and cdc42 lead to activation of MAPK pathways which further increase expression of CK2αP and one or more other proteins (?), and finically inhibit cell migration.

### Involvement of MAPK pathway

The mitogen-activated protein kinase (MAPK) signalling pathway is widely expressed in multicellular organisms, with critical roles in multiple biological processes, such as cell proliferation, death, differentiation, migration, and invasion. To understand the involvement of MAPK pathway in the OGR1 induced up-regulation of CK2αP, selective chemical inhibitors of Jun kinase (JNK), Erk and p38 were treated in the OGR1 or empty vector transfected A549 cells and transcript expression of CK2αP was analysed. The results showed that presence of selective inhibitors of JNK or p38 abrogated the OGR1 induced up-regulation of CK2αP expression whereas presence of selective inhibitor of Erk did not effect on the CK2αP expression upon OGR1 over-expression (**Figure 4B**). The overall results indicate that OGR1 inhibits A549 cell migration via up-regulation of CK2αP and NEP expression through activation of cdc42/Rac1 and MAPK pathways.

## Discussion

Although, OGR1 has been reported as novel metastasis suppressor gene, the molecular mechanism is yet to be understood. In this study, we demonstrate to show that OGR1 regulates expression of a cellular enzyme which acts as a regulator of several hallmarks of cancer cell behaviour CK2α [16–18] and an important membrane enzyme, neutral endopeptidase 24.11 (NEP, neprilysin, enkephalinase, CD 10). Our finding is the first report, to the best of our knowledge, to demonstrate that the expression of *CSNK2A3* (*CK2αP)* is regulated by a G protein coupled receptor, OGR1. Our results revealed that OGR1 up-regulates the expression *CSNK2A3* transcript but not the CSNK2A1 expression. CK2 is ubiquitously expressed and, in contrast it is believed to be a constitutively active enzyme and its regulation appears to be independent of known second messengers [51]. Considering the above findings, the result of immunoblotting against a specific CK2α antibody which shows up-regulation of CK2α protein expression when OGR1 is over-expressed in A549 cells indicates that the increased in CK2α protein may be the translated product of *CSNK2A3* (*CK2αP*). Therefore, our findings suggest that the aberrantly expressed transcript and/or protein of CK2α found in various cancer cells may be due to regulated *CSNK2A3* (*CK2αP*) expression which is potentially inducible or repressible by several master regulators of developmental pathways. The *CSNK2A3* might have an advantage over the CSNK2A1 in cancer cells in which sophisticated lineage specific genes expression is necessary [21]. However, further investigation is necessary to find out the expression of *CSNK2A1* and *CSNK2A3* in various cancer cells employing strategies which can distinguish expression from one another. It is important to mention that the mRNA sequence of *CSNK2A3* is 99.7% homologous to the *CSNK2A3* [21].

Our results further showed OGR1 also up-regulates expression of the transcript as well as protein of NEP, a cell surface cell surface peptidase that is normally expressed by numerous tissues, including prostate, kidney, intestine, endometrium, adrenal glands and lung. Loss or decreases in NEP expression have been reported in a variety of malignancies [52]. Reduced NEP may promote peptidemediated proliferation by allowing accumulation of higher peptide concentrations at the cell surface, and facilitate the development or progression of neoplasia. It has been shown that the effects of NEP are mediated by its ability to catalytically inactivate substrates such as bombesin and endothelin-1, but also through direct protein–protein interaction with other protein such as Lyn kinase [which associates with the p85 subunit of phosphatidylinositol 3-kinase (PI3-K) resulting in NEP-Lyn-PI3-K protein complex], ezrin/radixin/moesin (ERM) proteins, and the PTEN tumor suppressor protein [52]. To investigate whether CK2αP is upstream of NEP or vice-versa in the OGR1 signalling pathway, we analysed the protein expression of CK2α and NEP using immunoblotting in the presence of DL-thiorphan, a specific NEP inhibitor and silmitasertib (CX-4945), the specific inhibitor for CK2α respectively in the OGR1 over-expressed or control A549 cells. The result indicated that CK2αP is upstream of NEP in the signalling pathway of OGR1; indicating CK2αP induces NEP gene expression as result of OGR1’s action. The finding of CK2α’s regulation of NEP expression is of particular interest because of the role of CK2α in integrating to essential cellular processes such as cell growth, cell proliferation, cell survival, cell morphology, cell transformation and angiogenesis to proliferation and differentiation signals. In a variety of different cell types, peptide hormones including epidermal growth factor, insulin, and the NEP substrate bombesin increase CK2α activity. NEP plays a pivotal role in various cancers [40, 41]. It is previously reported CK2α inhibits NEP activity by phosphorylating cytosolic domain [49]. Therefore, our results reveal another layer of fine-tuning of NEP activity by regulating its expression induced by CK2αP, the key cellular enzyme.

In our previous study, we revealed that OGR1 inhibits prostate cancer cells migration via constitutive activity of Gαi [5]. To investigate whether OGR1’s up regulation of *CSNK2A3* in A549 is dependent of activation of Gαi, we analysed the transcript expression of *CSNK2A3* in the presence or absence of PTX upon OGR1 over-expression or empty vector. We used transcript expression of *CSNK2A1* as control. Our results revealed that OGR1 up-regulate expression of *CSNK2A3* transcript via Gαi activation.

To find out whether CK2αP and NEP are functionally involved in the OGR1’s induced inhibition of A549 cells, we assessed cell migration using would healing assay in the presence of specific casein kinase 2α inhibitor silmitasertib (CX-4945) or NEP specific inhibitor DL-Thiorphan following overexpression of OGR1 in A549 cells. The result revealed that inhibition of CK2α abrogates OGR1’s induced inhibition of A549 cell migration. The finding strongly suggested OGR1 inhibits cell migration via expression of CK2αP. Our finding supports the previous report that high levels of CK2αP correlate with higher survival rates of lung adenocarcinoma and renal clear cell carcinoma [32, 33]. The result also revealed that inhibitor of NEP abrogates the OGR1’s inhibition of A549 cells to some extent (30%). Our results also revealed that CK2αP is upstream of NEP and inhibition of casein kinase 2 abrogated completely (100%) the OGR1 induced inhibition of A549 cell migration whereas inhibition of NEP abrogated only 30%. Therefore, taken together our results suggest there may be other cellular molecule other than NEP to which CK2αP targets/activates following OGR1 activation.

To understand molecular mechanism of OGR1 induced up-regulation of CK2αP expression, we assessed roles of small G proteins; Rac1 & cdc42 and mitogen-activated protein kinases (MAPKs) pathways using negative dominant mutant of small GTPases and specific chemical inhibitors of MAPKs. The result revealed that inhibition of Rac1 and cdc42 activities by respective negative dominant mutant proteins completely abrogated OGR1 induced up-regulation of *CSNK2A3* whereas expression of *CSNK2A1* is not affected. Cells spread by putting out extensions that contact the surface, form adhesions, and then exert tension to induce outward movement. The process of contact the surface, form adhesions, and then exert tension to induce outward movement are reminiscent of the extensions and adhesions induced by the small GTP-binding proteins Rac and Cdc42 [53].

Further, we also assessed role of MAPKs; extracellular signal-regulated kinase (ERK), Jun kinase (JNK) and p38 since they have been shown to play a key role in transduction extracellular signals to cellular responses. The results clearly showed that inhibition of JNK and p38 completely abrogated OGR1 induced increase expression of *CSNK2A3* transcript but inhibition of ERK decreased *CSNK2A3 expression* some extend only. Taken together, these results indicated that OGR1regulates *CK2αP* expression via activation of small GTPase; Rac1 and cdc42 and MAPKs pathways.

The overall findings of this study clearly suggested that OGR1 regulates expression of *CK2αP* and *NEP* in A549 cells. OGR1 induced up-regulation of *CK2αP* through activation of small GTPase;α Rac1/cdc42 and JNK/p38 pathways are responsible for inhibition of cancer cell migration. However, OGR1 does not affect the expression of CSNK2A1. There is no previous report of how expression of CK2α in cancer cells is regulated although many studies have report of aberrant expression of the kinase in cancer. Therefore, our findings suggest that the aberrantly expressed transcript and/or protein of CK2α found in various cancer cells may be due to regulated CSNK2A3/CK2αP expression which is potentially inducible or repressible by several master regulators of developmental pathways. CSNK2A3 might have an advantage over the CSNK2A1 in cancer cells in which sophisticated lineage specific genes expression is necessary.

## Material and Methods

### Cell culture and reagents

The A549 cells (adenocarcinomic human alveolar basal epithelial cells) were procured from the NCCS, Pune, India. The cells were maintained at 37°C, 5% CO_2_ in F-12K Medium (Kaighn’s Modification of Ham’s F-12 Medium) supplement with 10% fetal bovine serum (FBS) and 1% penicillin/streptomycin. All the cell culture media, serum were procured from Gibco, USA while the antibiotics (penicillin/streptomycin) were procured from the Thermo Fisher Scientific (USA). The antibodies for ERK, phospho-ERK, JNK, phospho-JNK, p38 and phospho-p38 were purchased from the Cell Signaling, USA. Antibodies for Casein kinase 2 (Santa Cruz Biotechnology), OGR1 Cat No. 72500, abcam), NEP (Cat no. sc-9149, Santacruz Biotechnology) and β-actin (Cat no. sc-47778, Santa Cruz Biotechnology) were also procured. Specific inhibitors for JNK; SP600125, for p38; SB203580 inhibitors were procured from the abcam while the specific inhibitor for ERK, FR180204, silmitasertib (CX-4945), the specific inhibitor for Casein kinase 2α and DL-Thiorphan, the specific inhibitor for NEP were also procured from the Sigma-Aldrich.

### Plasmid construction and gene transfer

pcDNA3.1-OGR1 was constructed as described previously [5]. Briefly, the OGR1 coding sequence fragment (≈ 1.2 kb) was amplified and then cloned into pcDNA3.1 (puromycin) by *EcoR*I and *Hind*III digestion. The plasmid construct was transformed in DH5α competent E. coli, and selected colonies were then cultured. Plasmid DNA was purified using QuickLyse Miniprep Kit (Qiagen, USA). The construct containing OGR1 was further confirmed by sequencing using the ABI Prism 377 Automated DNA Sequencer (Applied Biosystems, Foster City, CA) and the DNA sequences and reading frames were further verified. The plasmids pcDNA3.1-cdcT17N and pcDNA3.1-RacT17N were procured from Addgene (Cambridge, USA). A549 cells were transiently transfected with 1.0–1.5μg plasmids; pcDNA3.1-OGR1 and empty vector (pcDNA3.1) or co-transfected empty vector or pcDNA3.1-OGR1 with pcDNA3.1-cdcT17N and pcDNA3.1-RacT17N using Lipofectamine 2000 (Thermo Fisher Scientific, USA) according to the manufacturer’s protocol. The final concentration of the SP600125, SB203580 and FR180204 was 20μM, and for silmitasertib (CX-4945) and DL-Thiorphan were 20μM, 2.5μM and 10μM respectively. All the inhibitors were added to the cells after 5 hrs of transfection. Treated cells were used for following experiments 48 hours after transfection.

### RNA isolation, cDNA synthesis and PCR amplification to measure the expression of casein kinase alpha and neutral endopeptidase

Total RNAs were extracted from 8×10^5^ cultured cells using RNA isolation kit (Qiagen, Germany) according to the manufacturer’s instruction. The isolated RNA were analysed for its integrity, purity and yield. Using the isolated RNA as template, first-strand complementary DNA (cDNA) was synthesised using M-MuLV Reverse Transcriptase (NEB, USA). Briefly, the extracted RNA (≈ 3μg) was reverse transcribed in a total volume of 20 μl with 350 μM dNTP, 50μM oligo(dT), 10X M-MuLV buffer, 200U RNase inhibitors and 200U M-MuLV reverse transcriptase (NEB). All the reagents were mixed and incubated at 42°C for 1 hour followed by 65°C for 20 minutes. PCR were then performed using platinum Taq DNA polymerase (TaKara, clontech; Japan). The reaction volume was 50 μl, containing 10X PCR buffer, 350 μM dNTP mixtures, 0.4 mM of each primer and 5U Taq DNA polymerase. Cycling conditions were 94°C for 4 minutes and then 25 cycles of (94°C 30s, 55°C 30s, and 72°C 45s) followed by 72°C for 10 minutes. Quantitative PCR were performed with primers obtained from Xcerlis genomics laboratory, Ahmedabad. The primer sequences for OGR1-specific primers; forward primers (*5′-CTGTCCTGCCAGGTGTGCGG-3*′ and reverse primers (*5′-CACGCGGTGCTGGTTCTCGT-3*’). The β2-microglobulin housekeeping gene (NM_004048) was used as the loading control. The primer sequences used to amplify the β-microglobulin were (forward) *5′- GAGCCTCGCCTTTGCCGATG-3*′ and (reverse) 5’CGATGCCGTGCTCGATGGGG-3’. The primer sequences for CSNK2A1 were (forward) *5′-CCAAACATCAAGTCCAGCTTTGTC-3′* and (reverse) *5′-ACCTCGGCCTAATTTTCGAACCA-3′* and for *CSNK2A3* were (forward) *5′-ATTGCTCCCCACTCCATCGC-3*′ and (reverse) *5′-ACCTCGGCCTAATTTTCGAACCA-3*′. The primer sequences for NEP were (forward) *5′-GCCTCTCGGTCCTTGTCCTGC-3*′and (reverse) *5′-ACGGGAGCTGGTCTCGGGAA-3*′.

### Cell lysate preparation and western blotting

Total cell lysates were prepared with radioimmunoprecipitation assay (RIPA) buffer. Briefly cells were lysed in 300 μl of RIPA buffer (10 mM Tris-HCl [pH 7.4], 150 mM NaCl, 1% Triton X-100, 5 mM EDTA, 1% sodium deoxycholate, 0.1% SDS, 1.2% aprotinin, 5 μM leupeptin, 4 μM antipain, 1 mM phenylmethylsulfonyl fluoride (PMSF), and 0.1 mM Na3VO4, sodium orthovanadate). Cell lysate was then centrifuge at 17,000xg for 1 hr and supernatant was mixed with 4X Laemmli sample buffer and equal quantity of protein samples were resolved by SDS-PAGE on 10% gels and transferred onto PVDF membranes. The membranes were blocked with 5% non-fat milk for 1 hr, incubated with primary antibodies (1:2000 dilution) at 4°C overnight and then with secondary antibody (1:6000 dilution) for 1hr at room temperature. The blots were detected using chemiluminescent ECL system (GE Healthcare). Blot’s Images were captured using ChemDoc (BioRad, USA).

### Wound healing assay

To analyse the effect of OGR1 and CK2α A549 cell was seeded in the six well plates so as to obtain 70-90% confluency at the time of transfection. After the serum starved for overnight, Cells were transfected with empty vector (pcDNA3.1) or pcDNA3.1-OGR1. After 5 hrs of transfection, cell-free area was created (scratched) in a 70–90% confluent monolayer cells using 10μl filter tips. The detached cells were removed by washing with 500 μL PBS. 2ml of fresh medium with or without Inhibitors; silmitasertib (CX-4945) and DL-Thiorphan were added afterwards and incubated. Cell migration was observed under the microscope and image was taken at 0-, 12-, 24- and 48- hrs).

## Acknowledgements

We are grateful to the Department of Biotechnology (DBT), Govt. of India and Science and Engineering Research Board, Department of Science and technology (DST-SERB), Govt of India for the research grant to LSS.

## Authors’ contributions

ALS, PMM & NTS performance of the experiments; LSS and ALS analyzed the data and wrote the first draft of manuscript; LSS & TRS reviewed & edited the manuscript; LSS supervised the project; LSS acquire the funding. All authors read and approved the final version of the manuscript.

